# A gene selection analysis of TAS1R1 and TAS1R3 to examine human carnivory

**DOI:** 10.1101/2023.10.17.562619

**Authors:** Mackenzie Cross, Andrew Kitchen

## Abstract

Humans are considered to have a unique reliance on meat compared to other primates, as much of humans unique evolutionary trajectory, such as human brain expansion, is linked to the increased consumption of meat for calories and nutrients. However, other primates such as chimpanzees (*Pan troglodytes*) and bonobos (*Pan paniscus*) are also known to consume meat. While humans meat consumption is considered to be unique in humans’ increased incorporation of tools to process the prey carcass and consumption of a broader range of prey, these distinctions are less obvious when contextualized within the broader behavioral repertoire of *Pan* species carnivory. This research seeks to identify if the taste perception of meat is different between humans and other ape clades through gene selection analyses. Specifically, this work examines the umami taste receptor genes, TAS1R1 and TAS1R3, which enable the savory flavor perceived when eating meat. Using PAML, we test for positive selection in these genes across several ape clades. We infer positive selection in TAS1R1 for the homininae clade and positive selection in TAS1R3 for the *Homo*/*Pan* clade. No selection was detected in only the human lineage, which complicates claims that human carnivory is unique compared to other primates while simultaneously suggesting the role of meat may be unappreciated in chimpanzees and bonobos as well as the role of insectivory in gorillas.

## INTRODUCTION

Humans consumed meat throughout their evolution (Braun, 2010; Dominguez-Rodrigo, 2005; Ferraro, 2013; Pickering, 2013; Semaw, 2003). There is evidence of stone-tool marks on bone in Gona, Ethiopia around 3.42-3.24 myr ago attributed to *Australopithecus afarensis* (McPherron, 2010), as well as faunal remains in direct association with Oldowan artifacts in the Ounda Gona South area of East Gona dated ∼2.6 Myr (Semaw, 2003), a large assemblage of cutmarked bones in Gona, Afar, Ethiopia dated at ∼2.6-2.5 Ma (Dominguez-Rodrigo, 2005), and later evidence of early transport and tool-assisted processing of bovid remains found at Kanjera South, Kenya at ∼2.0 Ma (Ferraro, 2013). Interestingly, this site features a prominent, yet unexpected quantity of medium-sized cranial elements, which Ferraro *et al*. suggest may be due to the nutrient dense, lipid rich qualities of these tissues overriding the energetic costs of their transport (Ferraro, 2013). Not only was meat consumed throughout hominin evolution (Braun, 2010; Dominguez-Rodrigo, 2005; Ferraro, 2013; Pickering, 2013; Semaw, 2003), but a broad range of prey were exploited, as there is evidence of both terrestrial and aquatic animal consumption as early as 1.95 Ma in East Turkana, Kenya (Braun, 2010). The frequency of humans’ meat consumption and their primary method of meat procurement is less clear. It is possible that hominins employed passive scavenging strategies, where hominins consumed the animal remains leftover from carnivore consumption, such as lipid rich bone marrow which may still remain after carnivore consumption and can be accessed via tool use (Blumenschine, 1991; Pobinar, 2020). This is in contrast to power scavenging where, rather than wait for the carnivore to leave the prey, hominins directly confront the carnivore to obtain earlier access to the prey (Pobinar, 2020). Indeed, more recent work indicates that hunting and/or power scavenging likely occurred more frequently (Domínguez-Rodrigo, 2003). Regardless, there is clear evidence linking human evolution to meat consumption.

Generally, meat is considered a “high quality” food source, as it is rich in both calories and nutrients. Specifically, meat consumption offers critical B-complex vitamins, essential amino acids, zinc, iron, selenium and phosphorus (Pereira and Vicente, 2013; De Smet and Vossen, 2016) in addition to protein and fat (Milton, 1999). Some of these nutrients are found predominantly in meat, while also deemed critical for human health, which some researchers infer as evidence for humans’ deep evolutionary ties to meat consumption. Iron is such a nutrient, as humans preferentially digest heam iron compounds obtained from meat, as opposed to plant based forms, which Henneberg et al. argue is an adaptation to meat eating (Henneberg, 1998). Furthermore, maintaining iron levels has been shown to be critically important for pregnant women and infants, as iron deficiencies can result in neurodevelopmental abnormalities (Lozoff and Georgieff, 2006). Other nutrients such as taurine, carnosine, and creatinine are found predominantly in meat and, simultaneously, are identified as critical to human health through their ability to inhibit oxidative stress (Wu, 2020). Although meat offers many nutrients and several of these nutrients serve important functions in human health, there are also negative impacts to a diet composed predominantly of protein consumption (Milton, 1999; Speth, 1988), as too much protein in humans negatively impacts health through increased stress to kidney and liver function, or results in micronutrient deficiencies, as there are many nutrients required for human health not present in meat (Speth, 1988). Carnivory encapsulates any type of prey consumption, thus including nutrient diverse muscle, viscera, bone marrow, organs, etc. (Milton, 1999). Indeed, the term meat eating has been criticized as too broad as it does not differentiate between these different parts of the carcass which contain vastly different nutrient profiles (Thompson, 2019). Beyond the region of the prey consumed, there is also great diversity in the nutrients offered by different species of prey, as marine and terrestrial prey can offer quite different nutrient profiles. For example, long-chain polyunsaturated fatty acids are found predominantly in marine prey. Interestingly, these fatty acids are also critical for normal human brain function (Broadhurst, 2002). Indeed, meat consumption, whether marine or terrestrial, offers critical nutrients often demanded by the human brain. A significant body of research links humans’ increased meat consumption to human brain evolution, as it is thought that meat provided some of the calories and nutrients required by the energetic demands of an expanding human brain (Pfefferle, 2011). Furthermore, Aiello and Wheeler suggested that a dietary transition to meat enabled the size of the intestines to shrink, thus releasing the intestines’ energetic demand to facilitate brain expansion (Aiello, 1995), although more recent research disputes this idea (Navarrete, 2011).

At least to some degree, human carnivory is acknowledged to hold importance to human evolution and human health. Although human carnivory is often discussed as distinct from other carnivorous primates, there are many parallels between human and non-human primate meat-consumption. Many primates across the order will opportunistically eat meat. Indeed, primates consume a range of prey species including groups such as invertebrates, amphibians, birds and other small mammals, as well as other primates (Watts, 2020). In particular, chimpanzees display quite diverse strategies in obtaining prey and exploiting the carcass. Chimpanzee tool use is well documented. Chimpanzees use tools to obtain prey, as well as more fully exploit the carcass. One of the first observations of chimpanzee tool-assisted hunting included a female chimpanzee from Mahale Mountains, Tanzania who used a modified branch to assist in capturing a squirrel (Huffman and Kalunde, 1993). Chimpanzees of the Fengoli savannah in Senegal habitually use tools to hunt. These chimpanzees construct a spear-like tool, which is used to remove bushbabies (*Galago senegalensis*) from tree cavities for consumption. Beyond tool use, chimpanzees also cooperatively hunt and share meat (Mitani, 2001; Watts, 2008). Furthermore, chimpanzees (*Pan troglodytes*) have been observed to scavenge, although this behavior is relatively rare (Watts, 2008). The carnivorous behavioral repertoire of closely related bonobos (*Pan paniscus*) is less established, although bonobos have been documented to prey on terrestrial ungulates and other primates (Hohmann and Fruth, 2007; Surbeck and Hohmann, 2008).

Given the relevance assigned to meat consumption during human evolution, it is important to clearly identify the aspects to human meat consumption which are distinct from other primates, specifically closely related chimpanzees and bonobos (hereafter, referred to as *Pan* species). Some work has explored humans’ relationship to meat through a genetic lens, such as scans for positive selection across many genes related to either protein and/or lipid digestion which appear to serve to make general claims across species (Babbit, 2011), but less of this work explicitly examines specific genes of interest. While there is a lot of work on examining various aspects of meat perception and digestion, this work is often contextualized within a medical context, as opposed to an evolutionary context. We propose the examination of genes critical to the taste perception of meat, as evidence of positive selection or lack thereof in these genes can assist in identifying which aspects of meat consumption serve special importance within the human lineage and may have contributed to humans’ unique evolutionary trajectory. Specifically, we present a gene selection analysis of the umami taste receptor genes, TAS1R1 and TAS1R3, to clarify the role of umami taste perception in humans and *Pan* species. The umami taste receptor enables the perception of “savory” flavor, which humans typically experience during consumption of meat, seafood, cheese and some vegetables, such as mushrooms and tomatoes (Chaudhari, 2009; Diepeveen, 2021; Wu, 2022). Specifically, this receptor responds to amino acids (Nelson, 2002) and proteinogenic molecules, such as monosodium glutamate (MSG) (Zhu, 2023) and can be enhanced when MSG is paired with 5’-nucleotide monophosphates, such as inosine 5’-monophosphate (IMP) and guanosine 5’-monophosphate (GMP) (Chaudhari, 2009; Wu, 2022). TAS1R1 and TAS1R3 genes encode dimer proteins that form the umami taste receptor. This receptor is thought to be the dominant receptor in the transduction of umami flavor, although there is some evidence which suggests other receptors may contribute to this perception as well (Chaudhari, 2009; Diepeveen, 2021; Zhu, 2023). While TAS1R1 and TAS1R3 have been examined for selection within humans (Valente, 2018), this is the first gene selection analysis exploring selection across primates.Although there are many facets to meat consumption in primates, this work specifically examines meat eating within the context of reliance on protein consumption.

## METHODS

DNA and amino acid sequence data of the human TAS1R1 gene (ENSG00000173662.21) and the TAS1R3 gene (ENSG00000169962.5) were downloaded from Ensembl. Additionally, all available primate orthologous sequences were downloaded for both genes, which included 22 total orthologues for TAS1R1 and 19 total orthologues for TAS1R3. To expand the sample size of both genes, the human DNA transcript from Ensembl for TAS1R1 and TAS1R3 were separately queried in NCBI’s BlastN using the standard database nucleotide collection for highly similar sequences. The DNA and amino acid sequences were downloaded from NCBI for all primates not included in the Ensembl data as well as the flying tree shrew (*Galeopterus variegatus*). In total, TAS1R1 was represented by 37 sequences and TAS1R3 was represented by 34 sequences. The Ensembl and NCBI sequences were combined into a shared file which was aligned using the amino acid sequences as a guide to maintain a proper reading frame of codons. Both TAS1R1 and TAS1R3 had a “trimmed” alignment, which removed insertions that less than 10 primates shared and a “selected” alignment, which removed species sequences which contained numerous gaps (sample size reduced to 34 in TAS1R1 and 33 in TAS1R3). An additional alignment version was generated for TAS1R3 termed “altered”, which manually adjusted two bases inconsistent with the amino acid reference, likely due to sequencing errors. Alignments for both genes underwent model tests to determine the most appropriate parameters for Phylogenetic Analysis Using PAUP* was used to generate the maximum likelihood tree. Both gene alignments underwent model tests to determine the most appropriate parameters for PAUP* input based on the data, as described in Swofford 1998 and Wilgenbusch, 2003. Thus, HKY85+G for TAS1R1 and GTR+G+I for TAS1R3 were used. Under these respective parameters, PAUP* (*Version 4*.*0a build 168 for ubuntu*) was used to generate a Maximum Likelihood tree (ML) and a consensus tree (500 bootstraps) for both TAS1R1 and TAS1R3. In the TAS1R1 ML tree, there were three polytomies, which are more than PAML permits by default. As these polytomies were distant from the groups of interest and predominantly impacted the cercopithecoidea clade, we chose not to force a bifurcation and instead recompiled PAML (*version 4*.*10*.*7 for Mac Os X*) to accommodate for a polytomy by changing MAXNSONS 3 to MAXNSONS 10 prior to analyzing TAS1R1 and TAS1R3. Codeml in PAML was used to test for positive selection on specified primate lineages through Codeml’s branch model (model=2, NSsites=0, CodonFreq=7, seqtype=1). We tested a human-only foreground branch, Homo/Pan clade foreground branch, Homininae (Homo/Pan/Gorilla) clade foreground branch and ape (Homo/Pan/Gorilla/Pongo/Hylobatidae) clade foreground branch. Both genes underwent four total branch model analyses to test for positive selection on each foreground branch. Both genes underwent a test for the null model (model=0, NSsites=0, CodonFreq=7, seqtype=1) (Álvarez-Carretero, 2023). All results were imported into R (*version 4*.*2*.*2 x86_64-apple-darwin1*.*0 64-bit*) where they underwent a likelihood ratio test to evaluate significance.

## RESULTS

Slightly different versions of the alignment underwent these tests to ensure quality of alignment for both genes and most results did not typically differ between these versions. TAS1R1 did not show selection in the human lineage, nor the ape and hominini clade. However, selection was detected in the homininae clade for both alignment versions (trimmed p-value = 0.0274; selected p-value = 0.0356). TAS1R3 did not show selection in the human lineage, nor the ape or homininae clade. Interestingly, positive selection was detected in the Homo/Pan clade that was significant across alignment versions (altered p-value = 0.010; selected p-value = 0.016) and highly significant in one alignment version (trimmed p-value = 0.0043). While results were largely consistent across alignment versions, the TAS1R3 homininae results were the only findings with conflicting results between alignments. One alignment produced a highly significant result (selected p-value = 0.017), while another neared significance (altered p-value = 0.084) and the last was not significant (trimmed p-value = 0.249). All other results were consistent across alignments in their significance.

## DISCUSSION

Humans did not experience positive selection unique to the human lineage on either TAS1R1 or TAS1R3, which is unexpected given the importance assigned to meat consumption during human evolution. The clades which experienced consistent positive selection, such as homininae in TAS1R1 and the *Pan*/*Homo* clade in TAS1R3, indicate that meat consumption, at least in terms of umami taste perception, may be more important than expected in African apes. Firstly, selection in the TAS1R1 gene in homininae, which includes gorillas, *Pan* species and humans, is quite unexpected, given that gorillas are not known to be carnivorous in the sense that they consume vertebrate prey. However, it is well established that gorilllas partake in insectivorous behaviors, such as predation on ants, termites and other insects (Deblauwe, 2003). Moreover, insects can feature as a prominent aspect of the gorilla diet. Two groups of gorillas in Bai Hokou, Central African Republic were documented to regularly incorporate insects with each group consuming insects over 34% and 83% of the days observed respectively (Cipolletta, 2005). While insectivory is often distinguished from other types of predation, insects also contain proteinogenic molecules and thus are also capable of triggering the same umami taste perception signally cascade as other types of prey. Thus, the detection of positive selection in homininae for TAS1R1 may reflect a deep evolutionary history for proteinogenic prey across members of this clade, although this may manifest in different types of predation. While Homininae also experienced selection in some TAS1R3 alignment versions, this result was not consistent, although it does demonstrate the need for more work to be done which considered gorilla insectivory within a broader predation framework. Finally, the Pan/Homo clade also experienced positive selection in TAS1R3, which is another intriguing, unexpected result. Selection in this clade, as opposed to selection in only humans, suggests that the taste perception of umami flavor, as encoded by TAS1R3, was selected upon in a common ancestor of this clade, thus this function serves special importance across humans and *Pan* species. While Pan species are known to consume meat, their meat consumption is often described as less frequent than humans and their differences in carnivorous behaviors are often highlighted more regularly than their similarities. Broadly, these results suggest that taste perception of umami flavor carries shared importances across two clades, homininae for TAS1R1 and Homo/Pan for TAS1R3, rather than unique importance to humans. This provides unique insights into the deep evolutionary history of predation, and how selection on genes related to the taste perception of proteinogenic molecules may have occurred much earlier than likely anticipated.

While these results provide exciting insights into possible larger trends of protein perception and digestion, there are important limitations to this work. First, TAS1R3 is pleiotropic and, although well known for its umami taste perception functionality, it encodes other functions as well. Specifically, TAS1R3 also forms a heterodimer with the TAS1R2 gene which results in a receptor that detects sweet flavor (Nelson, 2002). Due to the pleiotropic nature of TAS1R3, it is not possible to definitively say that it was selected for its umami taste perception function. TAS1R1 does not have alternative functions established within the literature, although this is possible. Beyond this, the functioning of the umami taste receptor gene may be different in other primates. There is some research that indicates that some primate umami taste receptors show variation in their response to specific molecules (Toda, 2021), which suggests these receptors may serve purposes beyond umami taste perception in other primates/ As a result, it is possible that these genes experienced selection in some primate lineages for purposes other than umami taste perception. The complex evolutionary history of these genes limits the full explanatory power of these analyses, and demands that further work is done across several genes related to protein taste perception and digestion to better contextualize this work. In sum, this research provides an interesting initial examination into the genes which underpin meat flavor perception, which show a surprising pattern that suggests protein focused meat-consumed, at least in terms of the TAS1R3, may serve shared importance in the Pan/Homo clade, rather than unique importance in the hominin lineage. Additionally, its detection of positive selection in the homininae clade for TAS1R1 suggests the that insectivory may be an especially important form of predation for this group, and that African apes may share a deep evolutionary history which includes a shared reliance on predation, whether through insectivory or canonical “meat eating” prey. The explanatory power of these results are limited by TAS1R3’s known pleiotropy and the complex evolutionary history of both genes. Thus, this work serves as an interesting introduction to the application of gene selection methods to ape carnivory, but would greatly benefit from increased research.

## Notes

### Competing Interest Statement

The authors have declared no competing interest.

